# Probabilities of multilocus genotypes in SIB recombinant inbred lines

**DOI:** 10.1101/697144

**Authors:** Kamel Jebreen, Marianyela Petrizzelli, Olivier C. Martin

## Abstract

Recombinant Inbred Lines (RILs) are obtained through generations of inbreeding until all alleles are fixed. In 1931 Haldane and Waddington published a landmark paper where they provided the probabilities of achieving any combination of alleles in 2-way RILs for 2 and 3 loci. In the case of SIB RILs where sisters and brothers are crossed at each generation, there has been no progress in treating 4 or more loci, a limitation we overcome here without much increase in complexity. In the general situation of *L* loci, the task is to determine 2^*L*^ probabilities, but we find that it is necessary to first calculate the 4^*L*^ “identical by descent” (IBD) probabilities that a RIL inherits at each locus its DNA from one of the four originating chromosomes. We show that these 4^*L*^ probabilities satisfy a system of linear equations that follow from self consistency. In the absence of genetic interference – crossovers arising independently –, the associated matrix can be written explicitly in terms of the recombination rates between the different loci. We provide the matrices for *L* up to 4 and also include a computer program to automatically generate the matrices for higher values of *L*. Furthermore, our framework can be generalized to recombination rates that are different in female and male meiosis which allows us to show that the Haldane and Waddington 2-locus formula is valid in that more subtle case if the meiotic recombination rate is taken as the average rate across female and male. Once the 4^*L*^ IBD probabilities are determined, the 2^*L*^ probabilities of RIL genotypes are obtained *via* summations of these quantities. *In fine*, our computer program allows to determine the probabilities of all the multilocus genotypes produced in such sibling-based RILs for *L* ≤ 10, a huge leap beyond the *L* = 3 restriction of Haldane and Waddington.

## 1 INTRODUCTION

There are numerous inference problems in population and quantitative genetics that require comparing experimental frequencies of genotypes to those expected “theoretically”. Examples include genetic mapping of genomic markers, localizing causal factors of diseases and quantitative traits, performing marker assisted selection etc (Lander & Schork (1994); Weir (1996); Walsh & Lynch (2018)). The *expected* frequencies of genotypes, hereafter referred to as probabilities, of interest in such studies often involve multiple loci (*Buckler et al.* (2009)) and are strongly dependent on population structure. In population genetics studies, the structure of *natural* populations is rarely perfectly known. That partly explains why, in both animal and plant genetics, controlled crosses are widely produced to ensure a specific population structure. Arranging the crosses to lead to homozygous lines is greatly advantageous as such lines can be reproduced “identically and indefinitely”. The simplest situation satisfying these criteria is that of *recombinant inbred lines* (RILs)(Crow (2007)) founded from two parents as displayed in Fig.1. Given two (generally homozygous) parents that are the founders of the RIL construction (*F*_0_), one first produces the associated hybrids (*F*_1_). Second, starting with these *F*_1_ individuals, one produces a sequence of generations *F*_2_, *F*_3_, etc by iterative inbreeding, crossing male and female siblings until formally at *F*_∞_ one reaches full homozygozity (fixation of the alleles at all loci). As seen in Fig.1, the genomes of the homozygous lines produced by this process are mosaics of the parental genomes.

**Figure 1:**
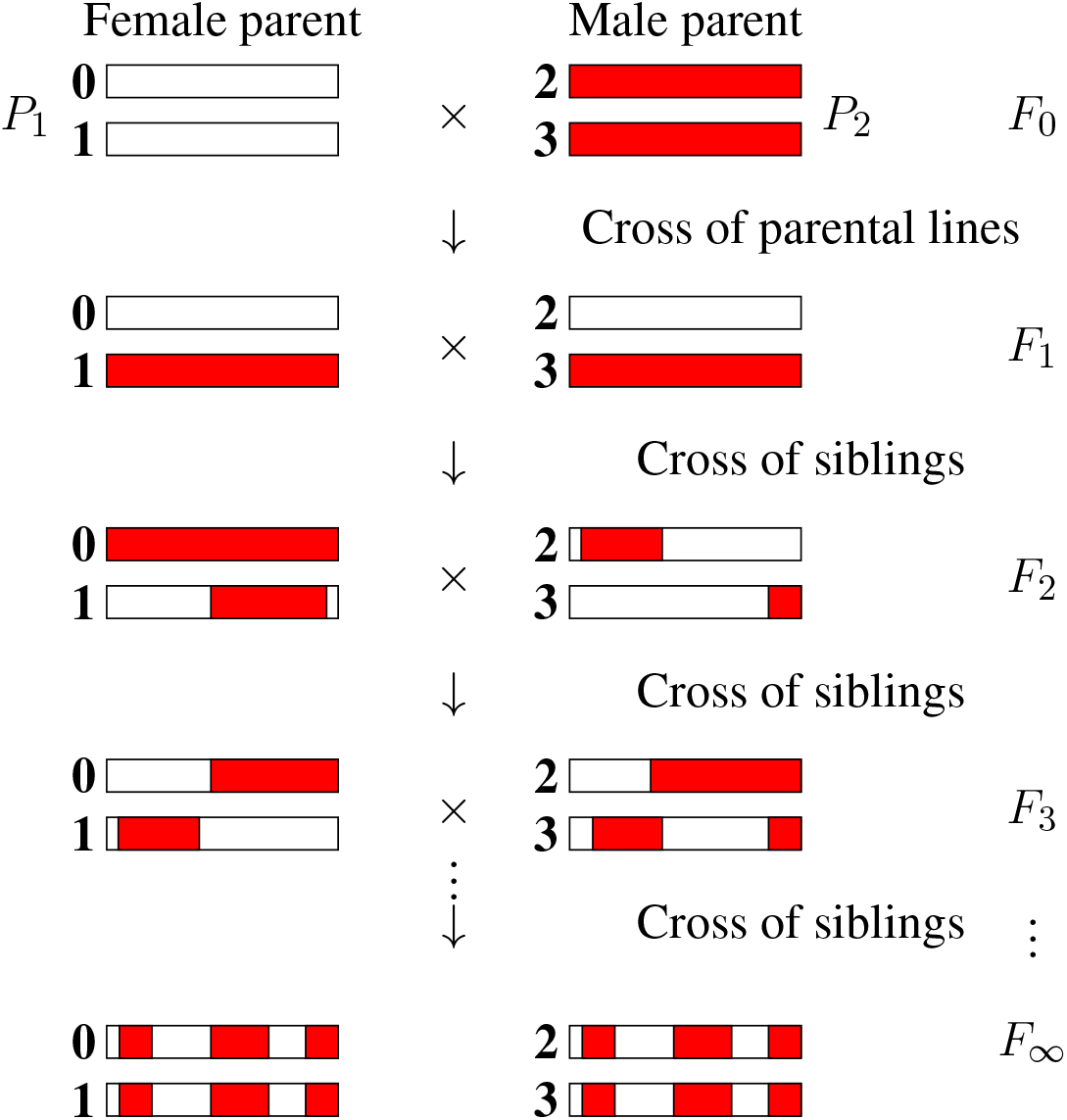
The production of recombinant inbred lines by sibling mating (SIB RILs). The homologous chromosomes (numbered 0, 1, 2, 3) inherit genomic segments from their parents, the boundaries of which identify crossover positions. The allelic content becomes fixed for “enough” generations, and so we introduce the limit of an infinite number of generations with the notation *F*_∞_.

Consider the allelic content at some set of *L* genomic markers or loci. There are then 2^*L*^ possible RIL genotypes, each having a probability that depends on how meioses generate recombinations between these different loci. In the case of plants that allow for selfing, the same individual is both the mother and the father of its offspring; the RILs are then produced *via* single seed descent (SSD) as opposed to *via* sibling (SIB) mating, this second case being the focus of the present work.

There are numerous generalizations of the RIL construction just given. Instead of using two parents to initiate the inbreeding, the use of 2^*k*^ parents leads to 2^*k*^-way RILs (Broman (2005)). 2^*k*^-way RILs start with 2^*k*^ parents to form 2^*k*−1^ offspring that are themselves crossed iteratively following a funnel (specifically a binary tree) pattern. Once the root of this tree is reached, the usual RIL inbreeding process is applied. For instance, the so called “Collaborative Cross” which has been a key community tool for mouse genetics, corresponds to *k* = 3; the choice there of using 8 founding parents at the top of the funnel allows for significantly greater allelic diversity than when using just 2-way RILs. Another generalization is the so called Advanced Intercross RIL (AI-RIL, sometimes referred to as Intermated RIL or IRIL) in which several generations of panmixia are inserted before applying the inbreeding to produce the RILs (Darvasi & Soller (1995); Rockman & Kruglyak (2008); *Winkler et al.* (2003)). Other generalizations include Multi-parent Advanced Generation Inter-Cross (MAGIC) (El-Din *El-Assal et al.* (2001)), *nested association mapping* (NAM) populations (*Buckler et al.* (2009)) etc. All of these population constructions involve some initial generations of allelic shuffling followed by the RIL (inbreeding) construction per se. Those *early* generations produce in effect initial conditions on the genotypes that are at the origin of the RILs and these initial conditions can be computed by direct recurrence from one generation to the next. In contrast, the RIL phase requires crossings that continue until all loci are homozygous and thus – at least mathematically – this phase involves an infinite number of generations. As a result, the computation of the probabilities of multilocus genotypes in RILs does not follow from a simple recursion over a fixed number of generations: either an extrapolation has to be made to deal with the infinite number of generations or some mathematical trick has to be devised to bypass the infinite nature of the process. This fact is at the heart of the difficulty of obtaining exact probabilities of multilocus genotypes in RILs.

The mathematical derivation of such RIL probabilities for two and three loci was provided by Haldane and Waddington (Haldane & Waddington (1931)) for bi-parental RILs in 1931. For two loci, by considering successive generations, they produced recursion equations for the probabilities of the corresponding (fixed or not) SIB genotypes which they then extrapolated to an infinite number of generations. This was quite a feat as they had to solve 22 simultaneous equations, leading *in fine* to their celebrated relation:

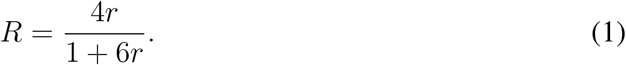

In this formula, *R* is the probability for a RIL two-locus genotype to be recombinant (have the allele of one *F*_0_ parent at one locus and the allele of the other at the other locus) while *r* is the recombination rate per meiosis between the two loci, assumed identical across male and female meiosis. We will rederive this formula using our framework in Section 3.1 because to our knowledge, the generalization of the Haldane-Waddington formula to situations where male and female recombination rates differ has not been published and our framework allows to deal with this extension.

Given *R*, it is easy to derive the probabilities of the four different RIL genotypes (each of the two loci can be fixed for either of the two parental alleles). Indeed, the two recombinant genotypes have the same probability and the sum of these two probabilities is precisely *R*. The probability of each of the two recombinant (respectively non-recombinant) RIL genotypes is then *R*/2 (respectively (1 − *R*)/2).

Haldane and Waddington further showed that this two-locus result also determined the three-locus probabilities. A way to see this is to notice that for three loci (*L* = 3) there are 2^*L*^ = 8 different RIL genotypes (at each locus the homozygous allelic state comes from one of the two parents). These 8 genotypes can be grouped into 4 pairs such that within each pair one genotype is obtained from the other by exchanging the alleles of the parents; for instance if the alleles of the parents are denoted by (A, B, C) and (a, b, c) at the three successive loci, the 4 pairs are {(*A, B, C*), (*a, b, c*)}, {(*A, B, c*), (*a, b, C*)}, {(*A, b, C*), (*a, B, c*)}, and {(*a, B, C*), (*A, b, c*)}. In each pair, the two complementary genotypes have the same probability so in effect it is enough to find the probabilities of each of the 4 pairs. These probabilities add up to one, providing a first equation. Then, labeling the loci as 1, 2, and 3, if the three meiotic recombination rates *r*_1,2_, *r*_2,3_ and *r*_1,3_ are known, the three RIL recombination rates *R*_1,2_, *R*_2,3_ and *R*_1,3_ are also. These quantities provide three further equations relating the four pair probabilities. These four equations uniquely determine the four pair probabilities and thus the probabilities of the 8 RIL genotypes.

Since that 1931 Haldane-Waddington landmark paper, some works have provided generalizations of Eq. 1, for instance in the case of 2^*k*^-way RILs (Broman (2005); Teuscher & Broman (2007)) and in the case of IRILs (*Winkler et al.* (2003); Teuscher & Broman (2007)). However, the problem of dealing with more than three loci seems substantially more difficult. Following the Haldane-Waddington algebraic approach, if there are *L* loci, there are 16^*L*^ possible allelic combinations at each generation and so it is necessary to diagonalize a 16^*L*^ × 16^*L*^ matrix; that task takes on the order of 16^3*L*^ operations and thus cannot be done on a standard computer even for *L* = 4. To our knowledge, the only work providing closed-form expressions for 4 or more loci is that of Samal and Martin (Samal & Martin (2015)) but their framework for determining exact probabilities of RIL multilocus genotypes applies only to single seed descent RILs, not to SIB RILs. The contribution of the present work is to show that the case of SIB RILs is also to a large extent tractable. In particular, (i) we give the analytic expressions for treating four loci in the absence of crossover interference, and (ii) we show that our framework allows to tackle more loci, though at a computational cost (CPU time and also computer memory) that increases roughly as 16^*L*^. Specifically, our computer scripts, written in R (Ihaka & Gentleman (1996)), can treat *L* = 8 loci in approximately 5 minutes when run on a desktop computer while a high-end server allows us to go up to *L* = 10 loci.

## 2 OVERVIEW OF THE METHOD

In the less complex case of single seed descent RILs, it was possible to determine the probabilities of the 2^*L*^ RIL multilocus genotypes by writing self-consistent equations directly associated with these unknowns (Samal & Martin (2015)). However, in the case of SIB RILs, the situation is more subtle because the allele carried by a RIL genotype may come from *either of the two siblings* at the *F*_1_ generation and thus “identical by descent” (IBD) does not reduce to identity by state (having the same allelic content) as can be seen in Fig. 1. As a result, it is necessary to first work with the 4^*L*^ probabilities that a RIL inherits IBD at the *L* loci from any of the four *F*_1_ homologous chromosomes. After introducing in Section 2.1 the 4^*L*^ RIL multilocus IBD probabilities, we show in Section 2.2 that each of these unknowns satisfies a self-consistent equation relating it to the others. These equations allow to overcome the technical obstacle of there being an unlimited number of generations in the process of generating RILs. Although these 4^*L*^ self-consistent equations constrain the 4^*L*^ unknowns, we show in Section 2.3 that one additional equation is necessary to specify the solution. For that last constraint we use the fact that the sum of all probabilities is 1. In Section 2.4 we show how the complexity of the problem can be reduced by working with a subset only of the unknowns. Finally, upon solving the system of equations to determine the IBD quantities, each of the 2^*L*^ RIL multilocus genotype probabilities follows by summing the probabilities of all compatible IBDs as will be shown in Section 2.5.

### 2.1 Probabilities of multilocus IBD inheritances in RILs and the set of non-equivalent *Q*’s

For a given RIL *L*-locus genotype (specified formally at generation *F*_∞_), the genomic content at any locus ℓ (ℓ ∈ {1,…, *L*}) will be IBD with exactly one of the four *F*_1_ homologous chromosomes. (One may note that the allelic fixation can happen before the IBD fixation, but no matter what, after an infinite number of generations both the IBD and the allelic states are fixed, that is they are identical across the four chromosomes of the SIB pair.) We number those four chromosomes 0, 1, 2 and 3 as indicated in Fig. 2 and use the same labeling for the later generations too. The IBD case illustrated is such that the RIL inherits from the *F*_1_ chromosome 2 at the first locus and from the *F*_1_ chromosome 1 at the second locus. (By convention we order the loci from left to right.) More generally, let us introduce the probability *Q*(*i*_1_, *i*_2_,…, *i*_*L*_) that a RIL inherits IBD from *F*_1_ chromosome *i*_ℓ_ for locus ℓ, ℓ = 1,…, *L* where *i*_ℓ_ = 0, 1, 2, 3. Naturally the sum of these 4^*L*^ probabilities (there are four possible values of *i*_ℓ_ at each locus ℓ) is equal to 1.

**Figure 2:**
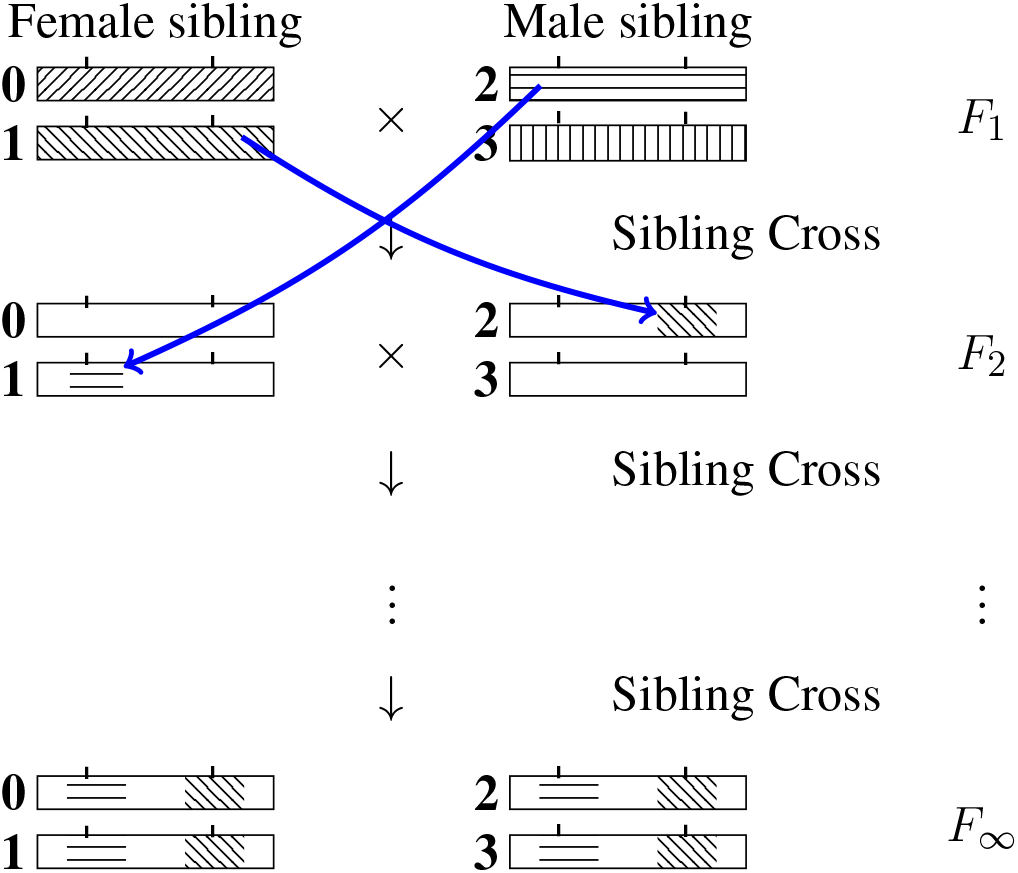
Inheritance during SIB mating and illustration of the construction of a self-consistent equation for any IBD probability. At each generation the homologous chromosomes are labeled 0, 1 (for the female) and 2, 3 (for the male). Note that the chromosomes labeled 0 and 2 are the outcomes of female meiosis while the chromosomes labeled 1 and 3 are the outcomes of male meiosis. The drawing illustrates the transition probability *T* [(2, 1) → (1, 2)] *Q*(1, 2) entering the self-consistent equation (*cf.* Eq. 3 when the left-hand side is *Q*(2, 1).

For *L* = 1, there are four IBD probabilities: *Q*(0), *Q*(1), *Q*(2) and *Q*(3). We shall assume Mendelian segregation with no bias in favor of any particular allele and so in particular the two homologues within each sex are equivalent. Then *Q*(*i*) = 1/4 for all *i* ∈ {0, 1, 2, 3}. Moving on to *L* = 2 for which there are 16 *Q*’s, the equivalence of homologues leads to the equalities *Q*(0, 0) = *Q*(1, 1), *Q*(0, 1) = *Q*(1, 0), *Q*(2, 2) = *Q*(3, 3), and *Q*(2, 3) = *Q*(3, 2) but also to equalities between mixed terms, *Q*(0, 2) = *Q*(0, 3), *Q*(1, 2) = *Q*(1, 3) etc. Furthermore, if female and male meiosis behave in the same way (so that in particular they have the same recombination rates), we can also conclude that *Q*(0, 0) = *Q*(2, 2) etc so that finally there are just *three* probabilities to determine, *Q*(0, 0), *Q*(0, 1) and *Q*(0, 2) instead of the initial 16. More generally, if there are *L* loci, how many *non-equivalent Q*’s are there? We shall assume there is no segregation bias and that female and male meioses have statistically identical behavior. Then it is possible to show (see Supplementary Material for details) that the number of non-equivalent *Q*’s is exactly

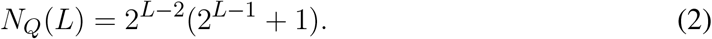

For example *L* = 1 leads to *N*_*Q*_(*L*) = 1 while *L* = 2 leads to *N*_*Q*_(*L*) = 3. The number of these non-equivalent *Q*’s grows roughly as (1/8) × 4^*L*^ to be compared with the total number ignoring equivalence of 4^*L*^. The factor (1/8) clearly makes it worth while to use such a reduction in the number of unknowns to simplify the task of writing and solving the equations. The proof of Eq. 2 in the Supplementary Material provides a way to enumerate the *Q*’s to be kept and schematically goes as follows. First, because all four chromosomes play equivalent roles, we can force *i*_1_ to be 0. Second, *i*_2_ can be constrained not to take the value 3 since that value can be replaced by 2, this time by equivalence of chromosomes 2 and 3. If *i*_2_ takes the value 0 or 1, we can again constrain *i*_3_ to be different from 3 by the same reasoning. If instead *i*_2_ = 2, then *i*_3_ must be allowed to take all values 0, 1, 2 and 3. We can proceed in this way to define the rules to be applied to the successive *i*_ℓ_. As long as the current list consist of 0s and 1s, the next *i* can be constrained to not take the value 3 by equivalence between chromosomes 2 and 3, but for all entries after the *first* occurrence of a 2, all values must be allowed (see the Supplementary Material for the final steps required to prove Eq. 2). As an illustration, the reader can check that for *L* = 3 loci, this construction leads to 10 non-equivalent *Q*’s, namely *Q*(0, 0, 0), *Q*(0, 0, 1), *Q*(0, 0, 2), *Q*(0, 1, 0), *Q*(0, 1, 1), *Q*(0, 1, 2), *Q*(0, 2, 0), *Q*(0, 2, 1), *Q*(0, 2, 2), and *Q*(0, 2, 3).

### 2.2 Self-consistent equations for the 4^*L*^ IBD probabilities

The IBD inheritance needs an infinite number of generations to become fixed with certainty, at least in principle. Our strategy consist in mapping such an infinite process into a finite one by relying on self-consistency. The probability for *F*_∞_ siblings to inherit IBD the sequence of “indices” (*i*_1_, *i*_2_,…, *i*_*L*_) from the *F*_1_ chromosomes can be decomposed into trajectories where the inheritance indices at the *F*_2_ level are also made explicit. If we denote these by 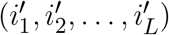, we can reinterpret *Q*(*i*_1_, *i*_2_,…, *i*_*L*_) as a sum of contributions:

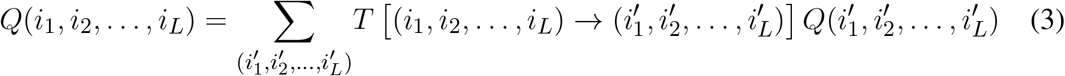

where *T* [(·) → (·)] is the transition probability of having the IBD propagate from the first list of indices to the second list of indices when going from the *F*_1_ to the *F*_2_ generation. *T* [(·) → (·)] is illustrated graphically in Fig. 2 by considering the case of two loci and having *i*_1_ = 2, *i*_2_ = 1, 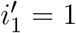 and 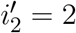.

Clearly *T* [(·) → (·)] depends on the meiotic process, and thus in particular on the recombination rates between loci. To simplify the notation, let us set *u* = (*i*_1_, *i*_2_,…, *i*_*L*_) and 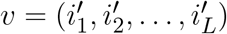. These transition probabilities *T* [(·) → (·)] satisfy three properties. First, if *i*_*k*_ = 0 or 1, *T* [*u* → *v*] = 0 unless 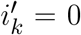 or 2. Similarly, if *i*_*k*_ = 2 or 3, *T* [*u* → *v*] = 0 unless 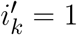 or 3. We summarize this via the rules

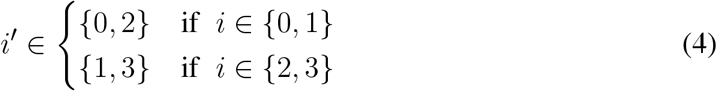

where *i* and *i*′ ∈ {0, 1, 2, 3}. Second, it turns out that the matrix *T* is “doubly stochastic” meaning that the sum of its entries in any row or in any column is exactly 1. The result that the sum over elements in a row is 1 follows from the fact that this sum gives the probability of having any of the possible outcomes of inheritances for a given starting point. Analogously, the result that the sum over all elements in a column is 1 corresponds to the fact that a given *v* is reached by some *u* and that summing over all possibilities for *u* again leads to 1. Third, each element of *T* decomposes into four factors,

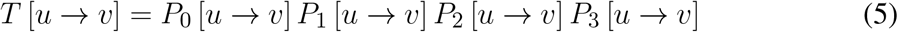

where the subscript of each *P* labels the chromosome of interest (and therefore the meiosis) at the *F*_2_ generation, thus *P*_*j*_ is a probability associated with the meiosis that produces chromosome *j* when going from *F*_1_ to *F*_2_. Consider for specificity the term *P*_3_. For the computation of this probability, only the entries in *v* equal to 3 matter. The corresponding *indices* specify which loci are thereafter IBD from chromosome 3 when considering the *F*_∞_ inheritance from the *F*_2_ generation. If those loci numbers are say 2, 5, and (*L* − 1), then *P*_3_ [*u* → *v*] is the probability for the loci 2, 5 and (*L* − 1) to inherit IBD from *i*_2_, *i*_5_ and *i*_*L*−1_ during the meiosis producing chromosome 3 when going from the *F*_1_ generation to the *F*_2_ generation. Note that all the other loci and chromosomes are irrelevant for this factor. The probability of that event is 0.5 (for the probability that the locus 2 will inherit IBD from chromosome *i*_2_) times the probability that the successive intervals 2-5 and 5-(*L* − 1) will be as required – recombinant or not – by the values of *i*_5_ and *i*_*L*−1_. Let us suppose that meioses arise without genetic interference, that is according to the so-called Haldane model (*Haldane et al.* (1919)). (Note that the values of these *P* s are the only part of our framework where crossover interference affects our computations; if these single-meiosis probabilities are known, then our framework provides the probabilities of all RIL multilocus genotypes just as in the case of no interference.) For specificity, if there is no interference and both intervals 2-5 and 5-(*L* − 1) are recombinant, the associated (meiotic) probability *P* is simply 0.5 × *r*_2,5_ × *r*_5,*L*−1_. Such a reasoning is easily extended to any situation, leading to the formula

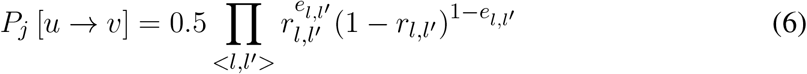

where the locus indices *l* and *l*′ are such that *v*_*l*_ = *v*_l′_ = *j*, *j* being the index appearing in the probability *P*_*j*_. In addition, the *e*_*l,l*′_ are defined as

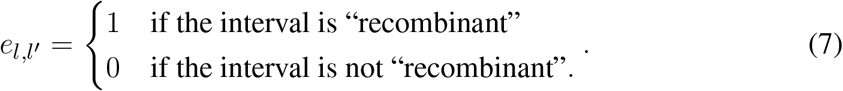

For Eq. 6, an interval *< l, l′ >* is called “recombinant” if and only if *i*_*l*_ and *i*_*l′*_ differ. Lastly, we need to specify the actual pairs of loci *l* and *l*′ that are to be used in that equation. To do so, we first construct the list of ordered indices that satisfy the constraint *v*_*l*_ = *v*_*l*′_ = *j*. The product in Eq. 6 is then over the *successive* pairs of this list. If the list is empty, *P*_*j*_ = 1 while if there is only one element in the list, *P*_*j*_ = 0.5. The interpretation of Eq. 6 is then as follows: there is a factor *r*_*l,l*′_ if the *u* list imposes that the interval *< l, l′ >* be recombinant and a factor 1 − *r*_*l,l*′_ otherwise. Putting together Eqs.3 and 5 specifies the 4^*L*^ linear homogeneous equations for the *Q*’s. In our computer software, we determine the matrix elements of *T* as formal mathematical functions of the *r*_*l,l*′_. In these general expressions it is possible to substitute the numerical values of the *r*_*l,l*′_ when necessary.

### 2.3 Adding one linear inhomogeneous equation to uniquely specify all 4^*L*^ IBD probabilities

Eq.3 can be rewritten as

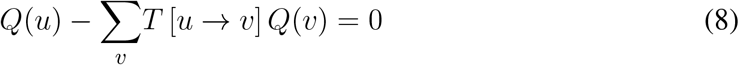

for all choices of *u*, corresponding to a set of 4^*L*^ linear homogeneous equations. Given one has as many equations as unknowns, one might hope that this system would determine the *Q*’s but that is not the case because these 4^*L*^ equations are not independent. Indeed, consider the sum of all the equations in the system:

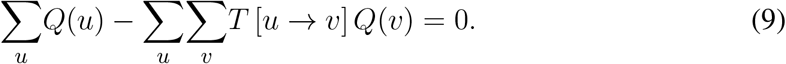

By interchanging the order of the sums this becomes

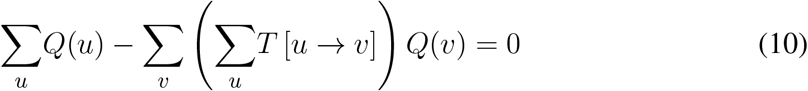

which is automatically satisfied because *T* is doubly stochastic so that 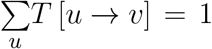.

To overcome the problem coming from this dependence amongst the homogeneous self-consistent equations, we need to include further information. We choose to do that by adding the constraint that the sum of all 4^*L*^ IBD probabilities equals 1:

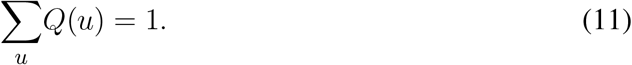

The inclusion of this (inhomogeneous) linear equation then uniquely specifies the values of all *Q*’s.

### 2.4 Reducing the system of equations to treat only the *N*_*Q*_(*L*) non-equivalent *Q*’s

As mentioned previously, it is advantageous to work with a subset of non-equivalent *Q*’s because this substantially reduces the complexity of the operations to be performed. Specifically, we modify the above approach by considering self-consistent equations only for the reduced list of unknowns – the *N*_*Q*_(*L*) non-equivalent *Q*’s chosen in Section 2.1 – so instead of having 4^*L*^ homogeneous equations of the type Eq. 8 we have only *N*_*Q*_(*L*) of them. In these *N*_*Q*_(*L*) equations, we replace each *Q*(*v*) by an equivalent *Q*(*v*′) where *Q*(*v*′) belongs to our list of *N*_*Q*_(*L*) unknowns. This recipe leads to *N*_*Q*_(*L*) linear homogeneous equations for our unknowns. Furthermore, we also apply these substitutions to the inhomogeneous equation Eq. 11, with the previously mentioned rule. As a result, by counting the number of *Q*’s arising in each equivalence class defined in Section 2.1, *Q*(*u*) occurs with weight 4 if the entries of *u* are all different from 2 and with weight 8 otherwise.

In practice, to solve this set of equations, it is convenient to have as many equations as unknowns so we remove exactly one of the homogeneous equations. In our computer algorithm we remove the last of these homogeneous equations but any other choice is just as valid. Having obtained as many independent equations as there are unknowns, the direct solution of this linear system (a linear algebra problem) provides the (unique) values of our *N*_*Q*_(*L*) non-equivalent *Q*’s.

### 2.5 Extracting the 2^*L*^ probabilities of RIL genotypes

Once the *Q*’s are determined, the probabilities of RIL multilocus genotypes can be computed by summing all IBD probabilities that are *compatible* with the RIL allelic content. Let us refer to the allelic content of parent 1 as a series of *A* alleles and that of parent 2 as a series of *a* alleles. Consider then a RIL multilocus genotype, written as a list *G* = (*α*_1_, *α*_2_,…, *α*_*L*_) of *L* alleles, *α*_*k*_ being *A* or *a*. The probability of a genotype *G* is obtained by summing over all *Q*(*u*) for which the *u* is compatible with the allelic content of *G*. The compatibility rule can be summarized as follows: if *α*_*k*_ = *A*, then *u*_*k*_ must be 0 or 2, while if *α*_*k*_ = *a*, then *u*_*k*_ must be 1 or 3. This is formalized mathematically by the following equation

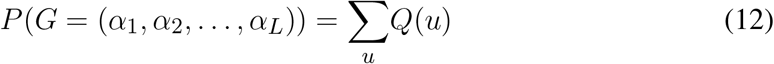

where the sum is restricted to the *u*’s satisfying the compatibility rule. Note that the *Q*’s on the right-hand side of Eq. 12 in general will not belong to our list of non-equivalent *Q*’s. As before, just omit all the terms associated with *Q*’s that are not in this list and multiply the other terms by either 8 or 4 depending on whether the associated *u* has one of its indices *u*_*k*_ equal to 2 or not, again because of the size of the equivalence classes.

## 3 RESULTS

We illustrate the power of our framework by considering increasing number of loci. The case of two loci is presented both for pedagogical reasons and to give the novel (as far as we know) values of the IBD probabilities when allowing for sex-dependent recombination rates. For three loci we detail the derivation of the coefficients of the self-consistent equations by giving associated graphical representations in the Supplementary Material. For four loci the analytical expression of the 40 × 40 matrix is also given explicitly. For more loci, the mathematical steps become too cumbersome to be dealt with by hand, but our computer code (in the form of R functions) can be used to first generate the analytic expressions for the linear system of equations, then to solve that system for the *Q*’s, and finally to produce the probabilities of all the RIL multilocus genotypes. The complexity of the computations provided by our framework can be summarized via the dimensionality of the linear system of equations used to compute the *Q*’s. This dimension increases roughly by a factor 4 for each additional locus for the simple reason that the number of unknowns increases in that way (*cf.* Eq. 2).

### 3.1 Case of two loci: recovering the Haldane-Waddington result and allowing for sex-dependent recombination rates

Haldane and Waddington (Haldane & Waddington (1931)) derived the formula for the probabilities of 2-locus RIL genotypes and Teuscher *et al*. (Teuscher & Broman (2007)) gave an alternative more compact approach. We will derive that Haldane-Waddington result here using our self-consistency approach. Then we show how to extend our framework to the case where female and male recombination rates differ.

Let *r*_*l,l*′_ = *r*_1,2_ denote the recombination fraction between the two loci (this recombination rate is for the moment taken to be the same in female and male as assumed by Haldane and Waddington). Furthermore, let *a*_*l*_ denote the allele at locus *l*, *l* ∈ {1,…, *L*}, on any of the homologous chromosomes in the RIL. By Eq. 2, for *L* = 2 there are 3 unknown *Q*’s. The indices *u* for each of these *Q*’s are such that they are not related by the symmetry between chromosomes. Our choice is to use *Q*(0, 0), *Q*(0, 1) and *Q*(0, 2). To build the 3 × 3 system of equations, begin with the inhomogeneous linear equation

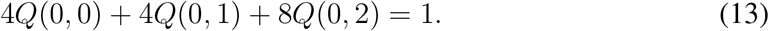

where the respective factors 8 and 4 follow from whether or not the *u* list of indices contains a 2. The next step is to write the self-consistent equation for each of the *N*_*Q*_(*L*) − 1 non-equivalent *Q*’s. For instance for *u* = (0, 0), by Eq. 3 applied to this case and using the rules for the vanishing of the elements of the matrix *T*, one has

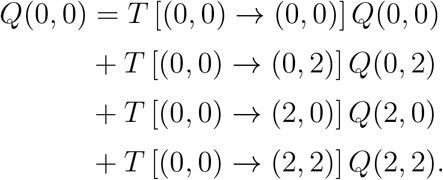

The matrix elements *T* [*u* → *v*] are determined by Eqs. 5 and 6. Direct calculation gives (1 − *r*_1,2_)/2, 1/4, 1/4, and (1 − *r*_1,2_)/2 respectively. To obtain a self-consistent equation involving only our three non-equivalent *Q*’s, we rewrite Eq. 14 by replacing *Q*(2, 0) by *Q*(0, 2) and *Q*(2, 2) by *Q*(0, 0), leading to

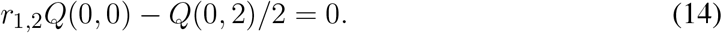

The self-consistent equation for *Q*(0, 1) is obtained by the same method. Eq. 13 together with Eq. 14 and its analogue for *Q*(0, 1) then lead to the system

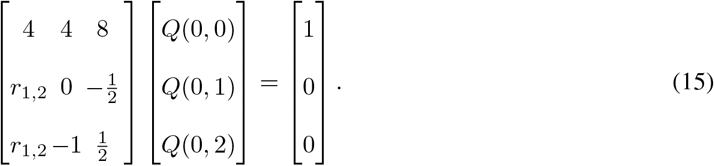

(Compared to Eq.8, we have changed the signs of each homogeneous equation to obtain a more readable matrix). This system can be solved by hand, leading to

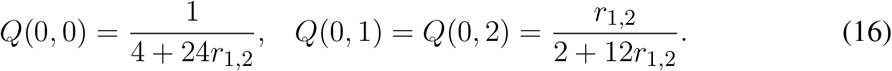

Given these three values, we can compute the RIL recombination rate *R* by summing all the probabilities of IBD events that produce recombinant RILs:

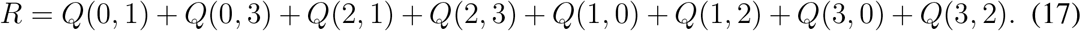

Using the equivalences (*Q*(3, 0) = *Q*(0, 2) etc), this gives *R* = 4*Q*(0, 1) + 4*Q*(0, 2); substituting the values from Eq. 16 leads directly to the Haldane-Waddington formula, Eq. 1.

How do these results extend to the case where female and male have different recombination rates, *r*^*f*^ and *r*^*m*^? The main complication comes from the fact that the symmetries of the system are reduced: one can no longer exchange the roles of female and male SIBs. As a result, there are 6 non-equivalent IBD probabilities. Without loss of generality, we take these to be *Q*(0, 0), *Q*(0, 1), *Q*(0, 2), *Q*(2, 0), *Q*(2, 2), and *Q*(2, 3). The determination of these six unknowns follows the same logic as when *r*^*f*^ = *r*^*m*^. First, use the inhomogeneous equation specifying that the *Q*’s are probabilities that add up to 1:

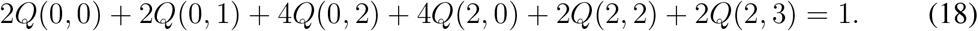

Second, determine the homogeneous equations associated with the self-consistency for the first *N*_*Q*_(*L*) − 1 non-equivalent *Q*’s. This then leads to the following system of equations:

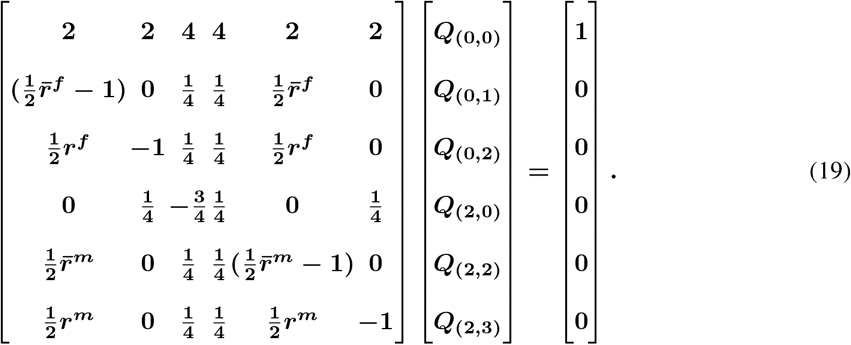

For the matrix elements in this system of equations, we have used the notation 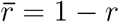 to designate the complementary value of the recombination rate, such a notation allowing for more compact expressions. The linear system Eq. 19 can be solved by hand, leading to

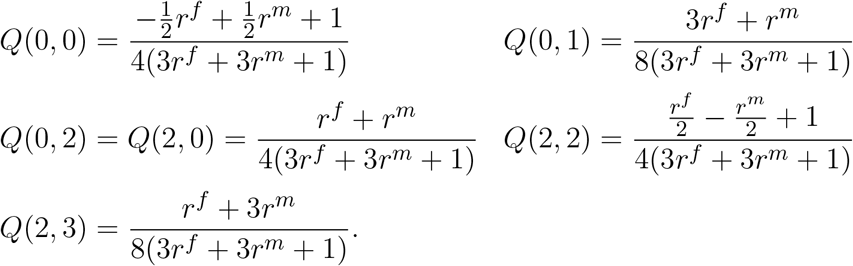

Note that except for *Q*(0, 2) and *Q*(2, 0), all the *Q*’s are *asymmetric* functions of *r*^*f*^ and *r*^*m*^. Furthermore, the equality *Q*(0, 2) = *Q*(2, 0) follows from the special symmetry of replacing the left-right convention that orients chromosomes by one using the right-left orientation.

Given the non-trivial result of Eq. 20, we can ask what is the consequence for *R*, the RIL recombination rate. The calculation is straightforward:

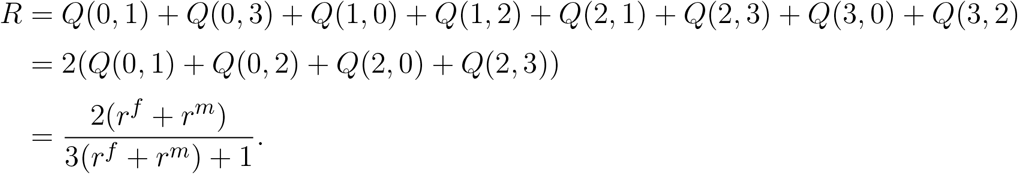

Interestingly, this result depends only on the mean of the female and male recombination rates, in spite of the fact that such a property does not hold at the level of the individual *Q*’s. Furthermore, it shows that the Haldane-Waddington relation (Eq. 1) can be used when recombination rates are sex-dependent if in that formula the (sex-independent) recombination rate is replaced by the sex-averaged recombination rate.

Although this example was very simple (it involved only two loci), it should be clear that our framework is generally applicable, for any number of loci, whether the female and male recombination rates are identical or not.

### 3.2 Case of three loci

Haldane and Waddington showed that the probabilities of two-locus RIL genotypes may be used to derive the probabilities of the three-locus RIL genotypes. Teuscher and Broman also provided this result when they introduced their approach (Teuscher & Broman (2007); Broman (2005)). In the introduction we explained why such a relation holds and so one might expect a similar conclusion to hold for the *Q*’s, but this is not so. Indeed, for this *L* = 3 case, as mentioned in Section 2.1, there are *N*_*Q*_(*L*) = 10 unknown *Q*’s to determine, corresponding to 9 degrees of freedom, but the information from the *L* = 2 level only provides 6 constraints, two for each pair of loci (6 = 2 × 3).

To determine the values of all the IBD probabilities, we simply apply our framework when using *L* = 3. We begin by specifying the set of non-equivalent *Q*’s that are our unknowns, following the logic of the general case as exposed in Section 2.1. We thus choose *Q*(0, 0, 0), *Q*(0, 0, 1), *Q*(0, 0, 2), *Q*(0, 1, 0), *Q*(0, 1, 1), *Q*(0, 1, 2), *Q*(0, 2, 0), *Q*(0, 2, 1), *Q*(0, 2, 2), and *Q*(0, 2, 3). Second, we write the single inhomogeneous equation that sums all *Q*’s (before applying equivalences). Third, we construct the self-consistent equations for the first 9 of our non-equivalent *Q*’s, assuming no genetic interference. The Supplementary Material provides a graphical representation of the *T* [*u* → *v*] entries to be explicit, our R code constructs this matrix automatically. These successive steps lead to the following linear system for our 10 unknowns:

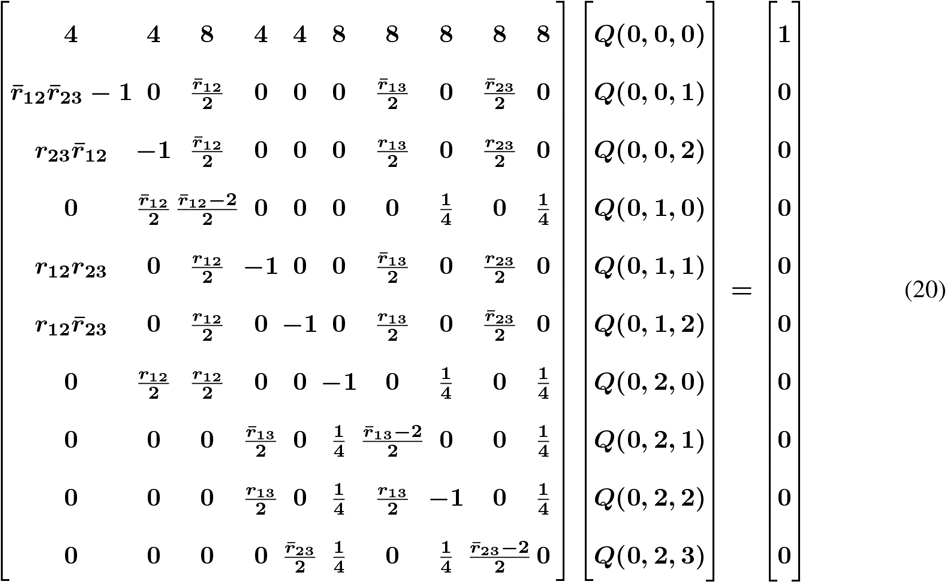

where as before 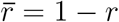 denotes the complementary value of the recombination rate and *r*_*ij*_ = *r*_*i,j*_. The solution of Eq. 20 can be obtained either numerically or analytically – that is as an explicit function of the three recombination rates – using e.g., Maple or Mathematica since a treatment by hand would be very tedious.

### 3.3 Four and more loci

The previous methodology can be extended to more loci but quickly becomes too cumbersome to manage manually. For illustration, in the case *L* = 4, there are 40 *Q*’s to determine (*cf.* Eq. 2). The system of 40 linear inhomogeneous equations determining these unknowns is given in Eq. 21 and barely fits on one page as a figure.

In that display including a 40 × 40 matrix, we have used the same compact notation as for *L* = 3. Our software produces this system of equations and then can solve for the *Q*’s for any particular values of the *r*_*ij*_. Computing the corresponding probabilities of RIL genotypes is then straightforward and in practice the computer does this very quickly.

It is of course possible to go to larger values of *L* but then it becomes unweildly to show the corresponding matrix. As expected, the computation time required by our R code grows fast with *L*, by about a factor 16 for each unit increase of *L*. The required computer memory also grows in the same way. At *L* = 8 the code takes about 5 minutes to solve the problem, and for still larger values of *L* it is best to use a server with large memory capacity (we have gone up to *L* = 10).

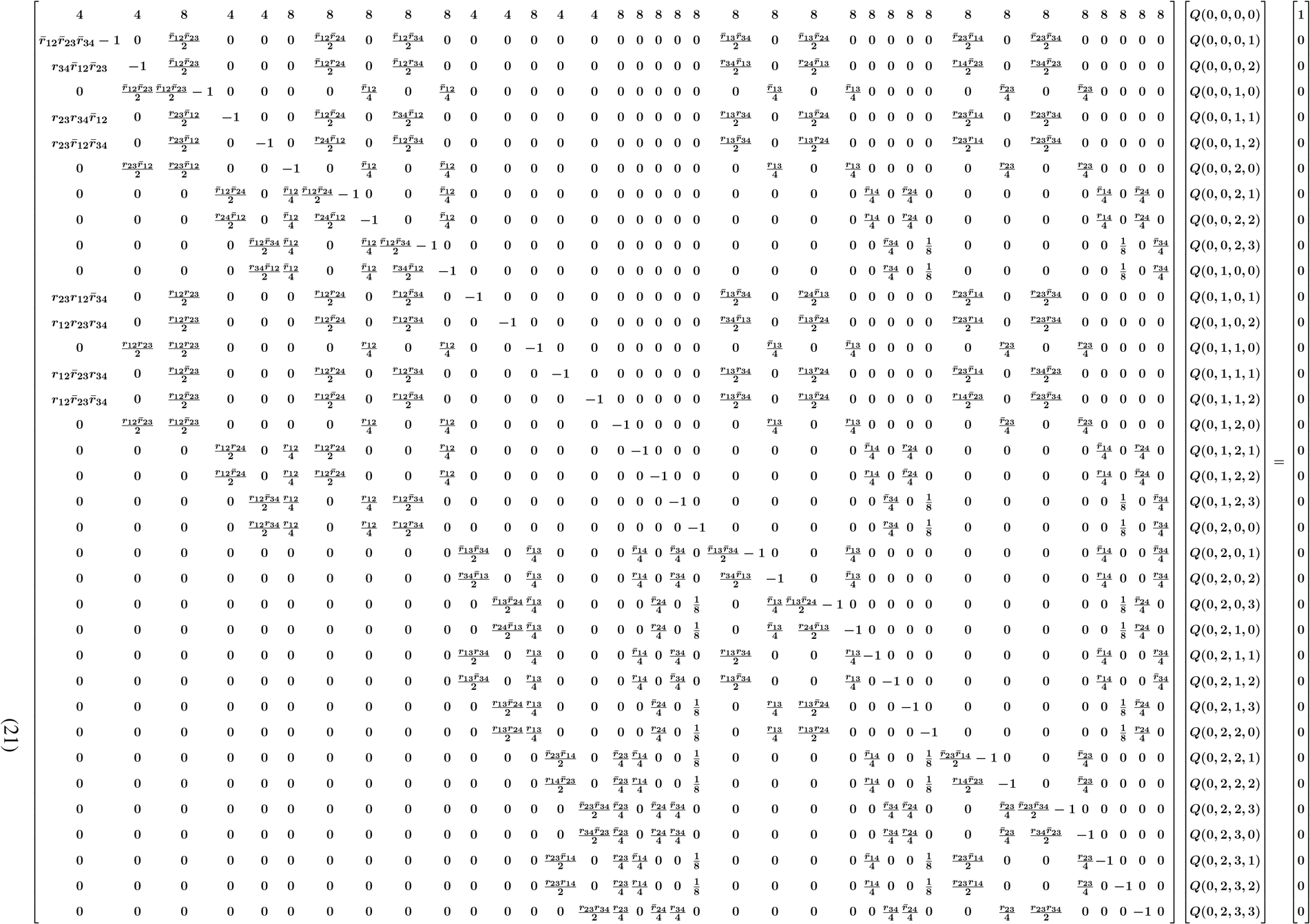

## 4 DISCUSSION

The construction of RILs involves successive generations of inbreeding until all alleles are fixed. The probabilities of the multilocus genotypes encountered across successive generations can be followed by recursion equations (Haldane & Waddington (1931); *Hospital et al.* (1996)) which in the case of SIB mating are specified by a dense matrix of size 16^*L*^ × 16^*L*^ if one has *L* loci. For the goal of obtaining the probabilities of RIL genotypes (the expected frequency of occurrence when averaging over a large number of repeats), the difficulty is that fixation formally requires an infinite number of generations. Thus, either the recursions must be taken “sufficiently far” to obtain *numerical* convergence or a mathematical trick has to be found. For *L* = 2, Haldane and Waddington succeeded in the second path thanks to much mathematical ingenuity, and interestingly, that *L* = 2 solution automatically determines the probabilities in the *L* = 3 case. However, since that founding work – going back to 1931 – no solution had been proposed to tackle SIB RILs with *L* = 4 or more.

Using a novel method, we have successfully overcome that long-standing challenge here. Our approach provides an algebraic solution, albeit at a computational cost that grows roughly as 16^*L*^ for *L* loci. That exponential growth rate is far less drastic than that of the original proposition of Haldane and Waddington of 1931, so that not only did we break the *L* = 4 barrier but in fact we were able to rather easily treat *L*’s up to 8. We also pointed out that our framework can deal with different female and male recombination rates, a situation that seems to have never been considered before in the context of SIB RILs, even for *L* = 2.

The ability to compute probabilities of RIL multilocus genotypes opens up to a number of applications. For instance, when building genetic maps, the ordering of markers is determined by comparing likelihoods of different orderings. That calculation can now be done using exact rather than approximate multilocus genotype frequencies, putting those mapping algorithms on a more solid footing. Similarly, when RIL genotypes must be inferred because of missing data, determining the most likely value of an allele requires comparing multilocus genotype probabilities. Finally, beyond specific uses in the case of RILs, our framework that exploits self-consistency might be useful in certain population genetics problems involving an infinite number of generations.

## AUTHORS CONTRIBUTIONS

OM proposed the project and with MP conceived and implemented a first approach. KJ introduced the analytic formulation and this led to major enhancements to the algorithmic KJ and MP developed the R scripts and all authors wrote, edited and approved the manuscript.

## ACKNOWLEDGMENT

The authors are grateful to Prof. D. de Vienne and C. Dillmann for insightful comments.

## SOFTWARE AVAILABILITY

R code implementing the methodology described in this paper is available online at https://github.com/olivier-c-martin/PMG_SIB_RILs.git

## SUPPLEMENTARY MATERIAL

The supplementary material contains two parts: a mathematical proof of Eq.2 from the main text, and the graphical representations of the self-consistent equations for the *L* = 3 case. The Supplementary Material for this article can be found online at: https://github.com/olivier-c-martin/PMG_SIB_RILs.git.

## Supplementary Material

### 1 SUPPLEMENTARY MATHEMATICS

Here we prove the equation

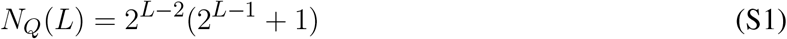

**Proof:** If *L* is the number of loci, there are 4^*L*^ IBD (identical by descent) probabilities *Q*(*i*_1_, *i*_2_,… *i*_*L*_) where *i*_*l*_ = 0, 1, 2 or 3 and furthermore these probabilities add up to 1. A number of these probabilities are equal because of two symmetries: (1) the two homologous chromosomes in each individual play identical roles, and (2) the siblings play identical roles (assuming no sex-dependence of meiosis, so that the recombination rates *r*_*l,l*′_ are sex-independent). It is thus appropriate to use only one representative of each equivalence class generated by these symmetries. A way to do this is to first impose that this representative have its first index, *i*_1_, equal to zero. Second, we can then specify exactly one element in each class by imposing that the indices of the representative *Q*’s have either

1. *i*_*l*_ ∈ {0, 1} ∀*l* ∈ {2,.., *L*}, *or*
2. *i*_*l*_ ∈ {0, 1} ∀*l* ∈ {2,.., *K* − 1}, *i*_*K*_ = 2 and *i*_*l*_ ∈ {0, 1, 2, 3} ∀*l* ∈ {*K* + 1,.., *L*}

The number of equivalence classes and thus of *Q*’s to consider is then

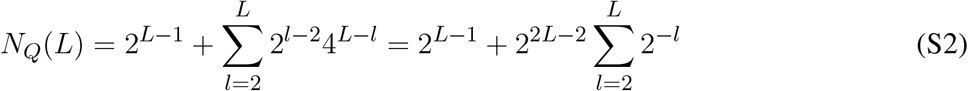

Given that 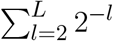 is a geometric progression of common ratio 2^−1^ from 2 to *L*, the sum of its terms can be expressed as:

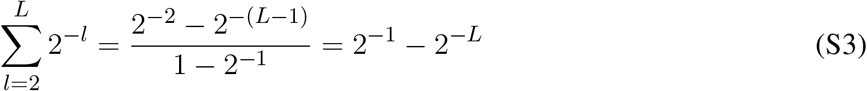

Substituting S3 in S2, we get

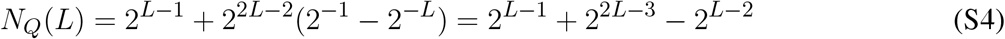

Factorizing with respect to 2^*L*−2^ and after simplification, this gives

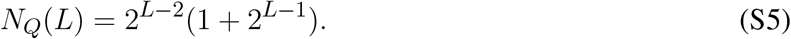

## 2 THE SELF-CONSISTENT EQUATIONS FOR THREE LOCI

Here we provide the coefficients entering each of the *N*_*Q*_(*L*) = 10 self-consistent equations for *L* = 3.

### 2.1 The self consistent equation for *Q*(0, 0, 0)

Figure S1 displays the 8 factors in the self-consistent equation for *Q*(0, 0, 0):

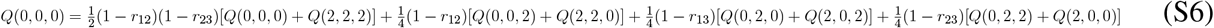

After use of symmetry to keep only non-equivalent *Q*s, this leads to

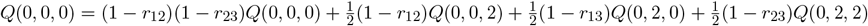

**Figure S1:**
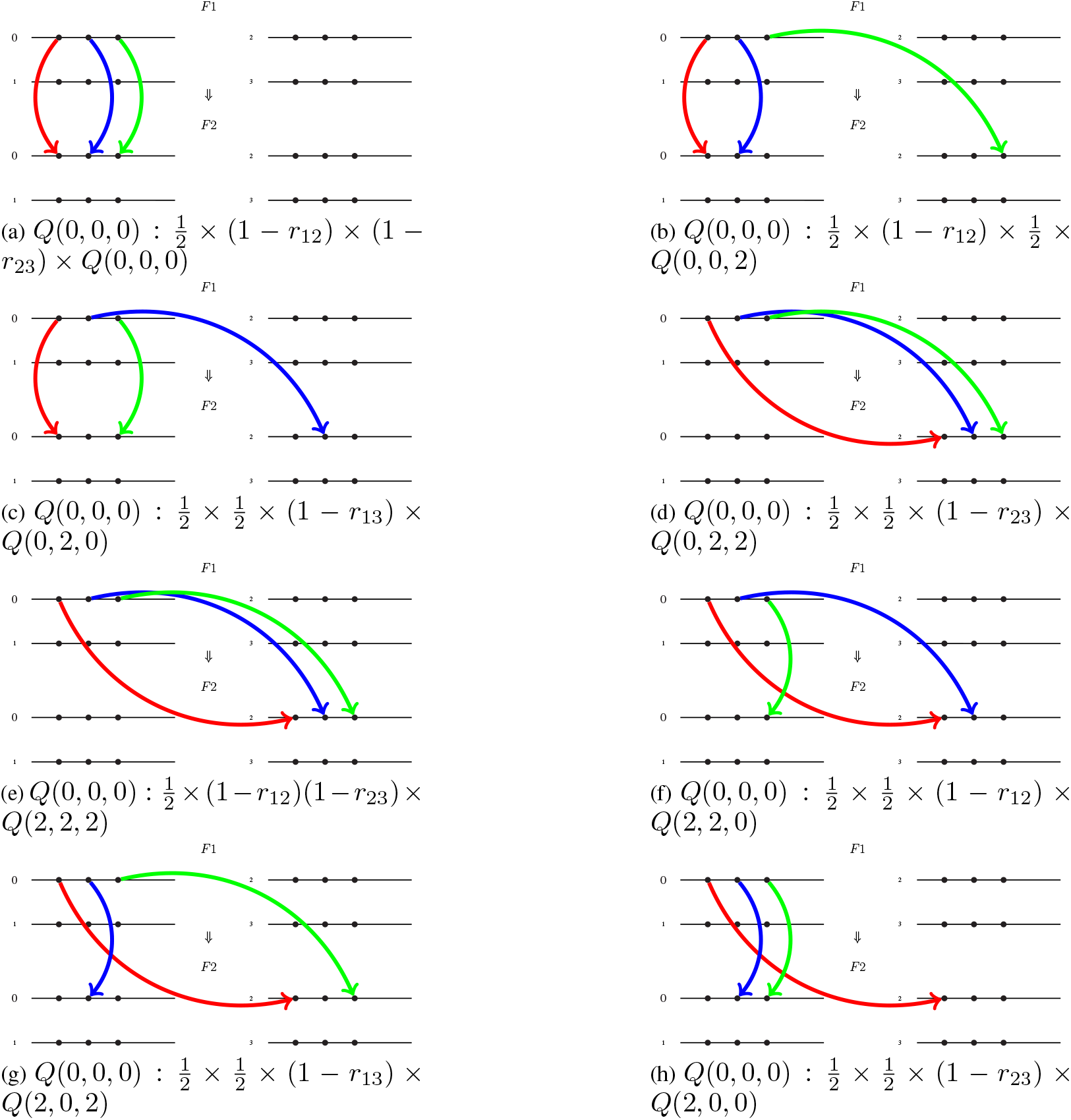
The graphical representation of the factors multiplying each *Q* on the right-hand side of Eq. S6 for *Q*(0, 0, 0).

### 2.2 The self consistent equation for *Q*(0, 0, 1)

Figure S2 displays the 8 factors in the self-consistent equation for *Q*(0, 0, 1):

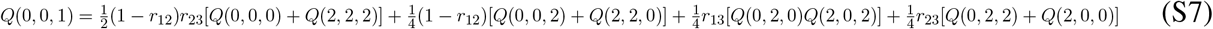

After use of symmetry to keep only non-equivalent *Q*s, this leads to

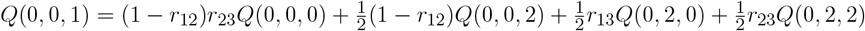

**Figure S2:**
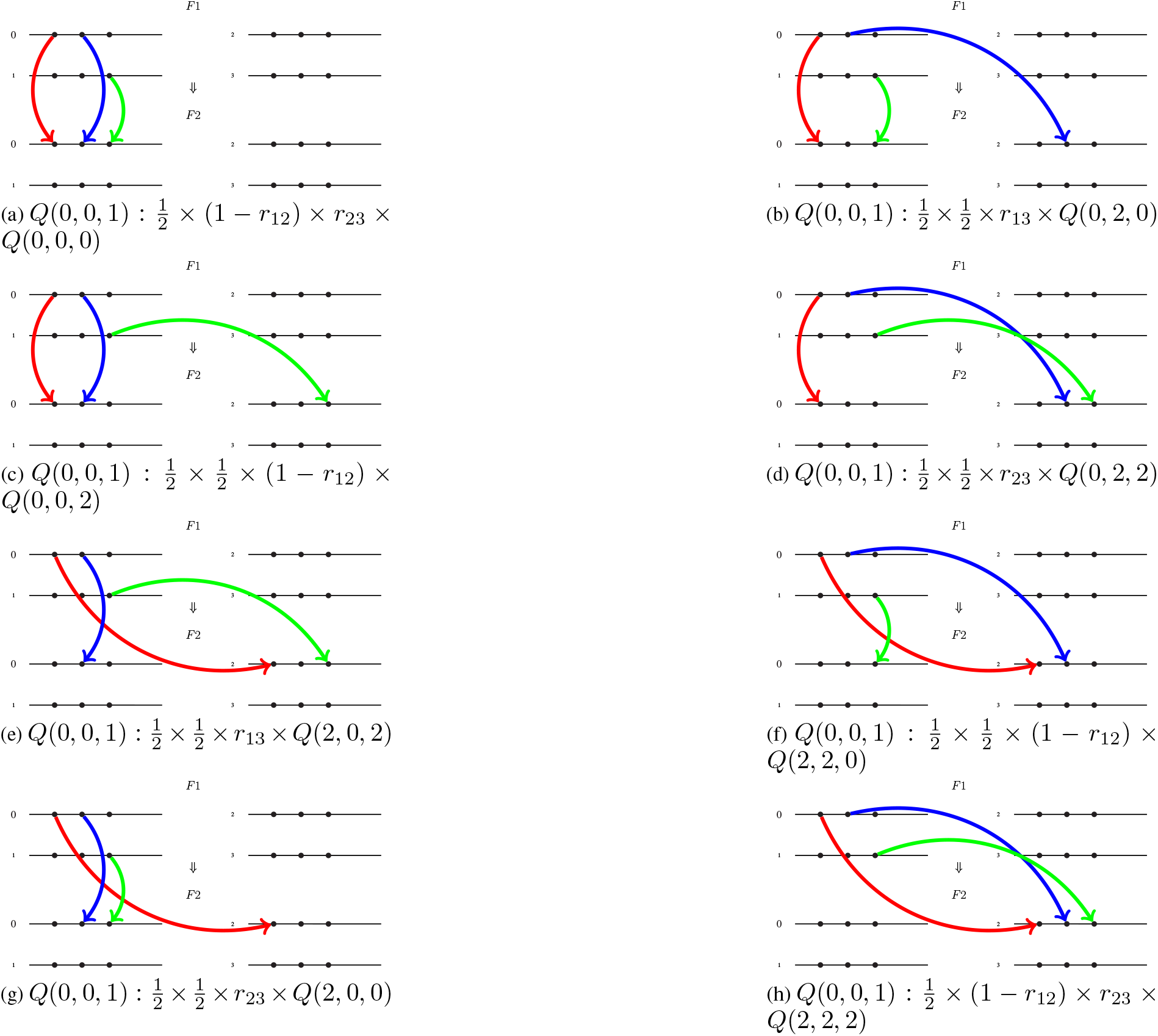
The graphical representation of the factors multiplying each *Q* on the right-hand side of Eq. S7 for *Q*(0, 0, 1).

### 2.3 The self consistent equation for *Q*(0, 0, 2)

Figure S3 displays the 8 factors in the self-consistent equation for *Q*(0, 0, 2):

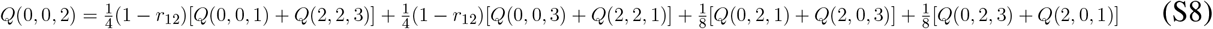

After use of symmetry to keep only non-equivalent *Q*s, this leads to

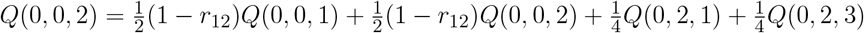

**Figure S3:**
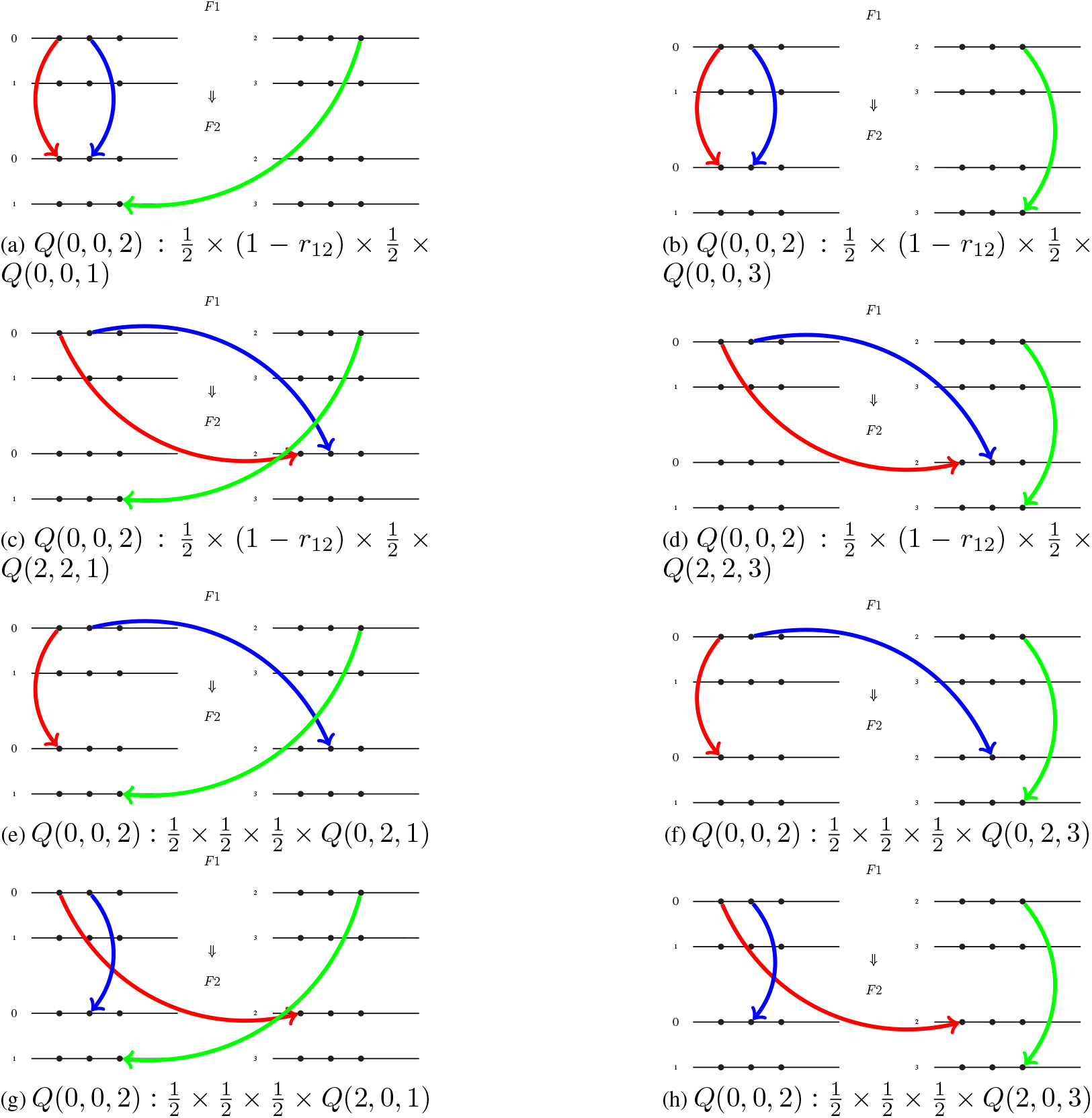
The graphical representation of the factors multiplying each *Q* on the right-hand side of Eq. S8 for *Q*(0, 0, 2).

### 2.4 The self consistent equation for *Q*(0, 1, 0)

Figure S4 displays the 8 factors in the self-consistent equation for *Q*(0, 1, 0):

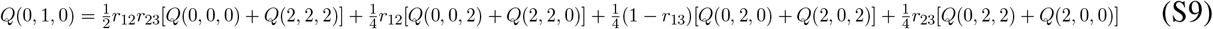

After use of symmetry to keep only non-equivalent *Q*s, this leads to

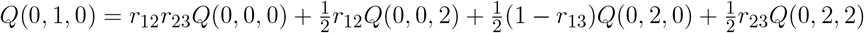

**Figure S4:**
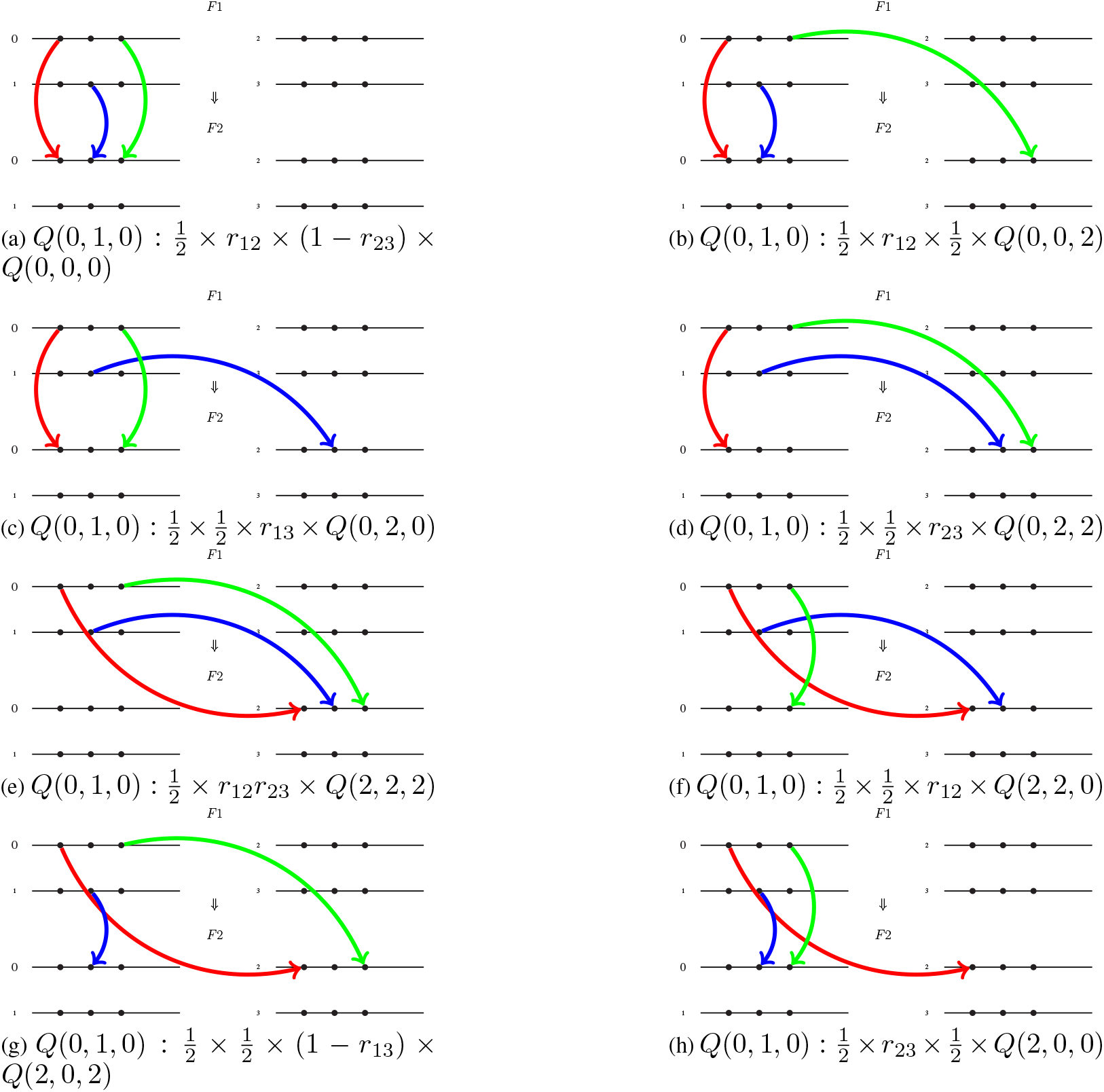
The graphical representation of the factors multiplying each *Q* on the right-hand side of Eq. S9 for *Q*(0, 1, 0).

### 2.5 The self consistent equation for *Q*(0, 1, 1)

Figure S5 displays the 8 factors in the self-consistent equation for *Q*(0, 1, 1):

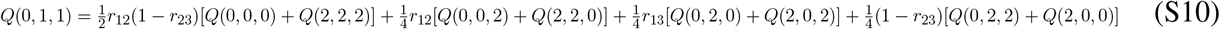

After use of symmetry to keep only non-equivalent *Q*s, this leads to

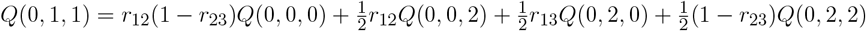

**Figure S5:**
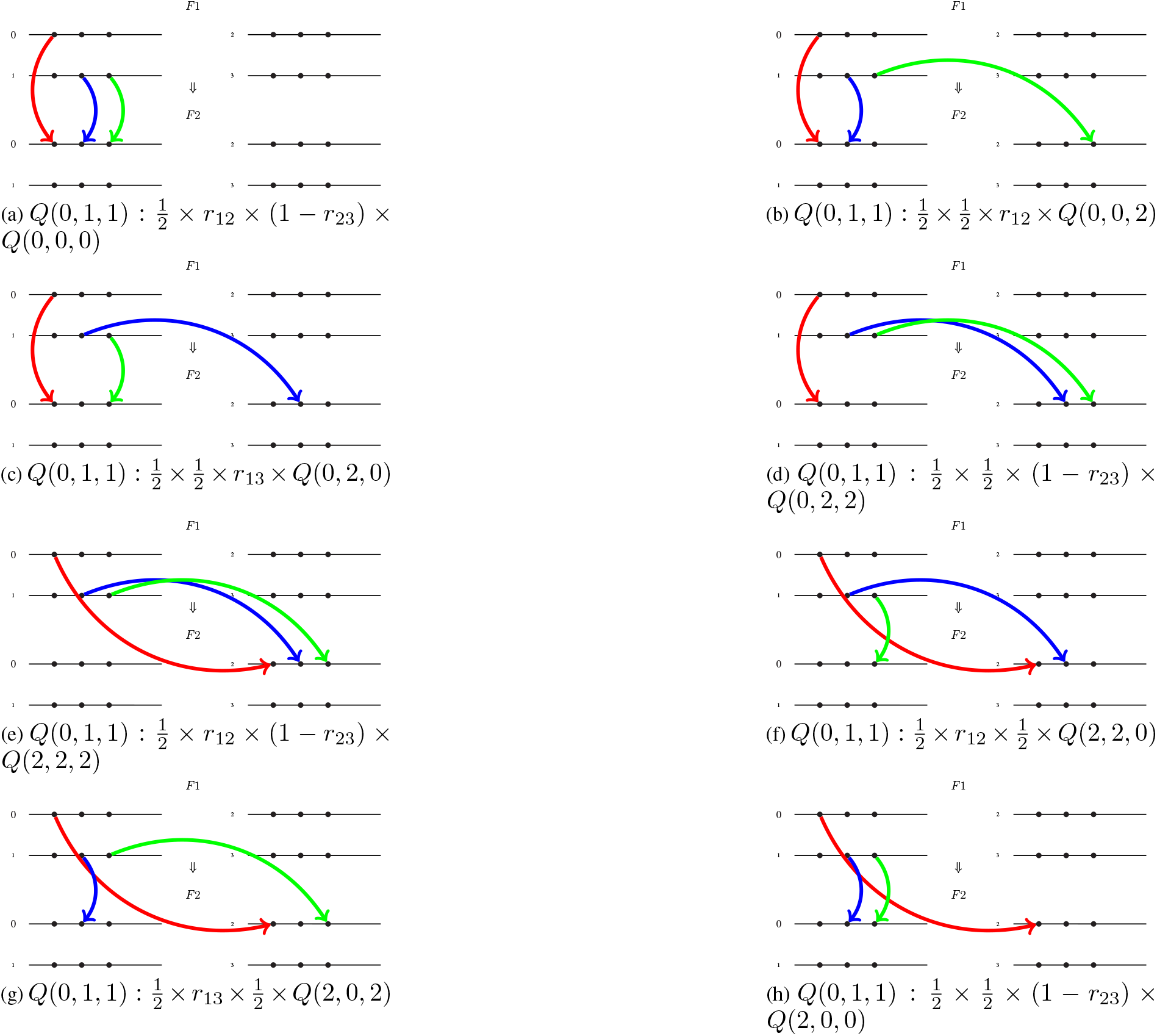
The graphical representation of the factors multiplying each *Q* on the right-hand side of Eq. S10 for *Q*(0, 1, 1).

### 2.6 The self consistent equation for *Q*(0, 1, 2)

Figure S6 displays the 8 factors in the self-consistent equation for *Q*(0, 1, 2):

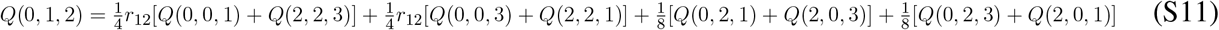

After use of symmetry to keep only non-equivalent *Q*s, this leads to

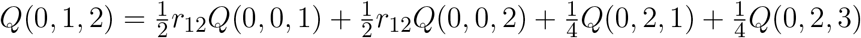

**Figure S6:**
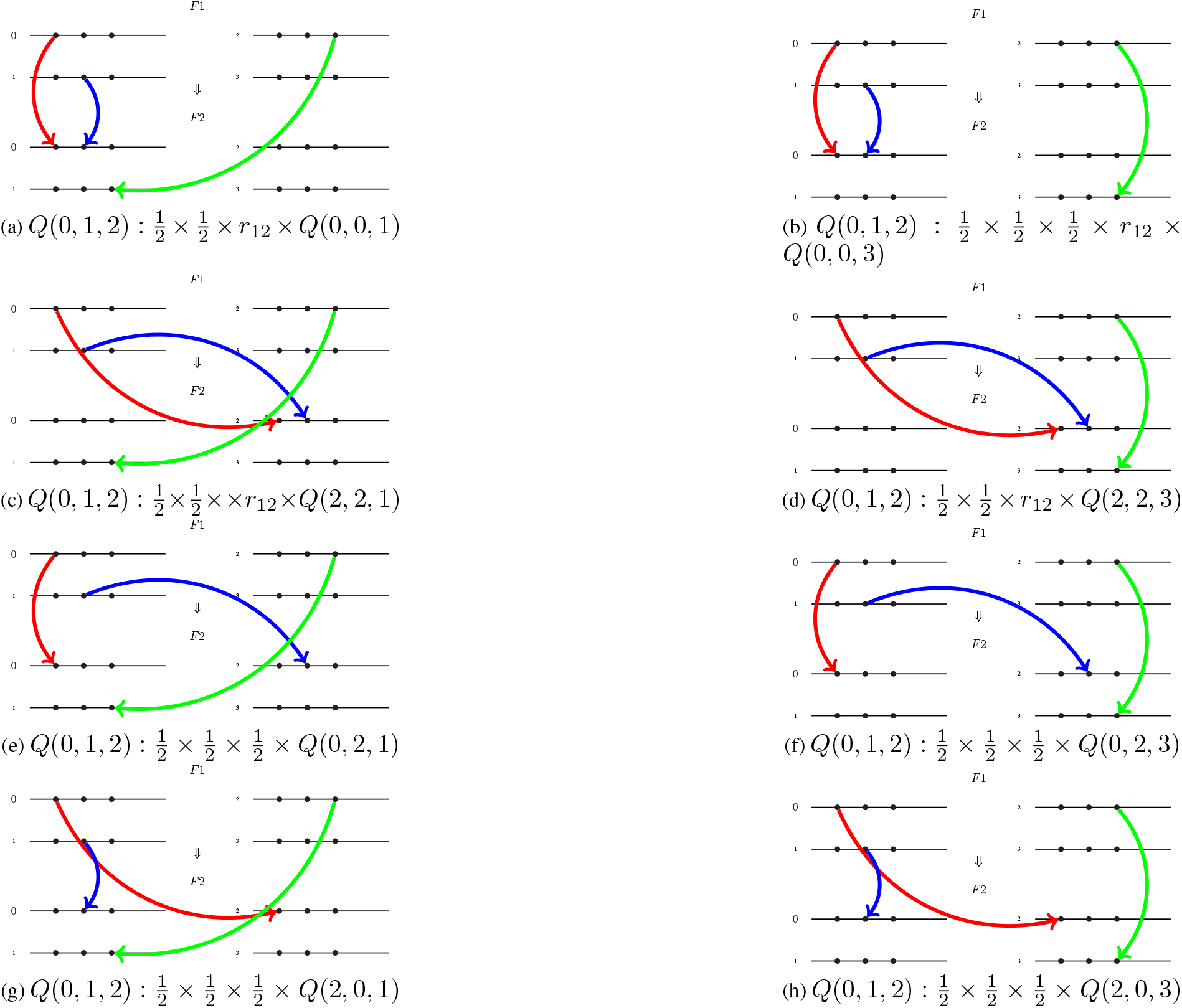
The graphical representation of the factors multiplying each *Q* on the right-hand side of Eq. S11 for *Q*(0, 1, 2).

### 2.7 The self consistent equation for *Q*(0, 2, 0)

Figure S7 displays the 8 factors in the self-consistent equation for *Q*(0, 2, 0):

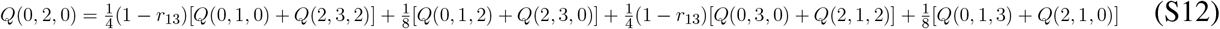

After use of symmetry to keep only non-equivalent *Q*s, this leads to

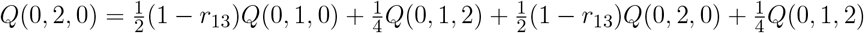

**Figure S7:**
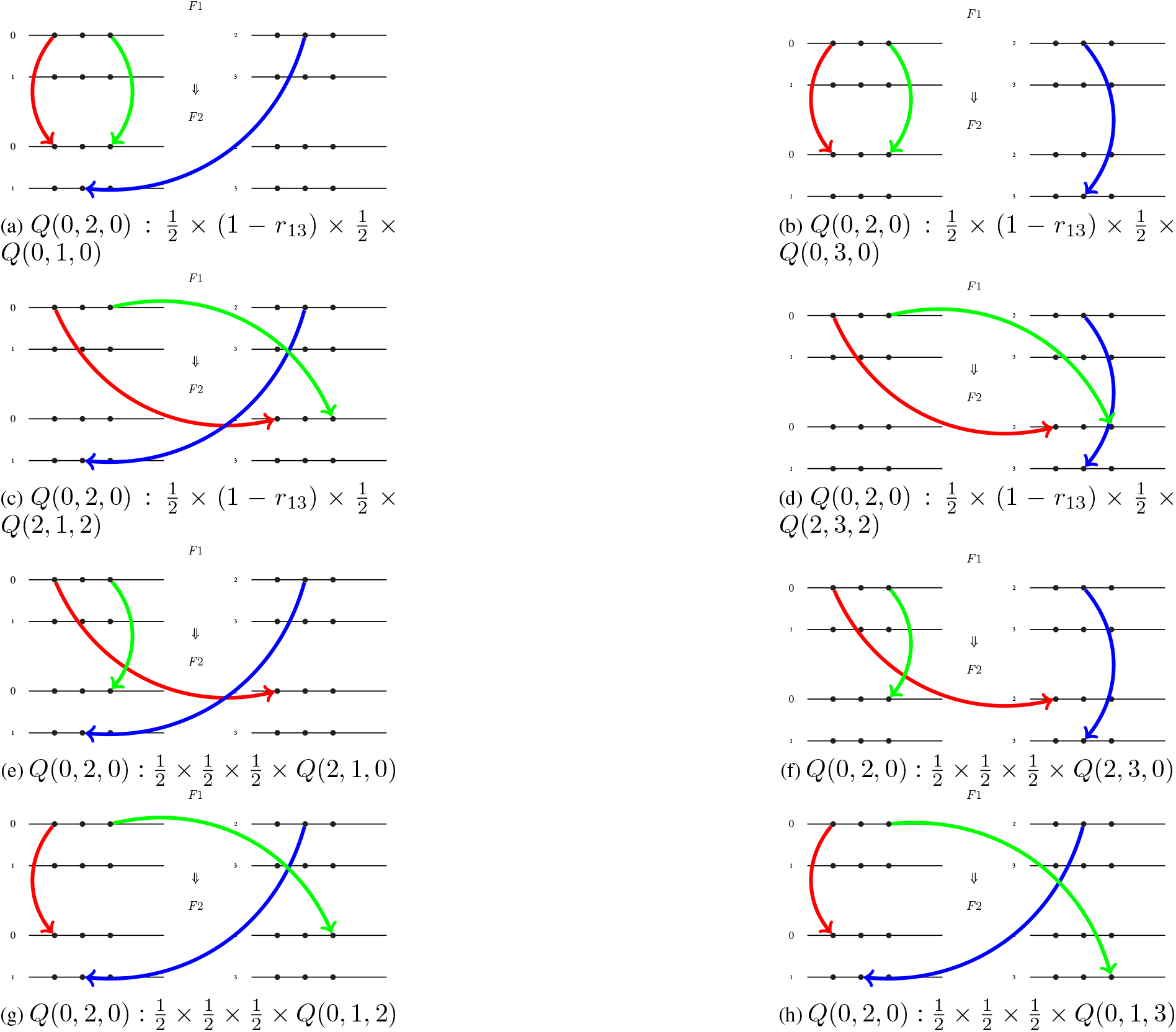
The graphical representation of the factors multiplying each *Q* on the right-hand side of Eq. S12 for *Q*(0, 2, 0).

### 2.8 The self consistent equation for *Q*(0, 2, 1)

Figure S8 displays the 8 factors in the self-consistent equation for *Q*(0, 2, 1):

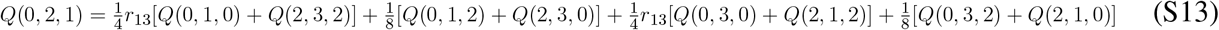

After use of symmetry to keep only non-equivalent *Q*s, this leads to

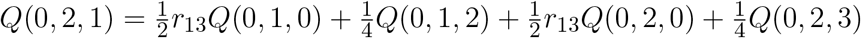

**Figure S8:**
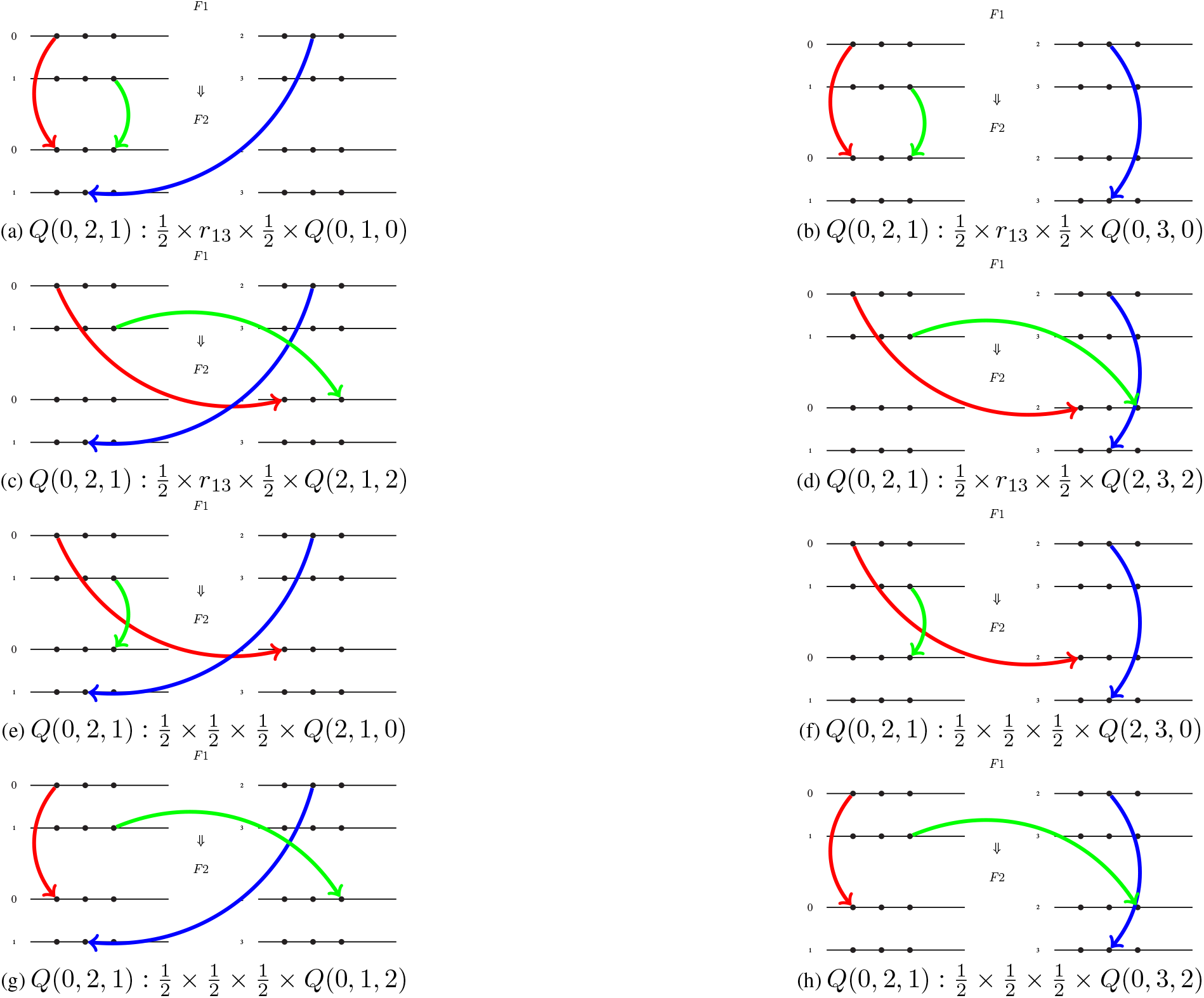
The graphical representation of the factors multiplying each *Q* on the right-hand side of Eq. S13 for *Q*(0, 2, 1).

### 2.9 The self consistent equation for *Q*(0, 2, 2)

Figure S9 displays the 8 factors in the self-consistent equation for *Q*(0, 2, 2):

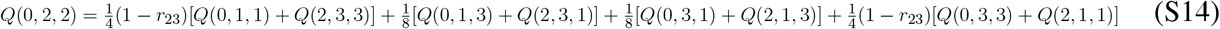

After use of symmetry to keep only non-equivalent *Q*s, this leads to

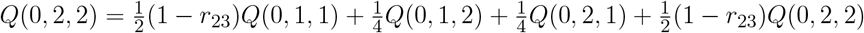

**Figure S9:**
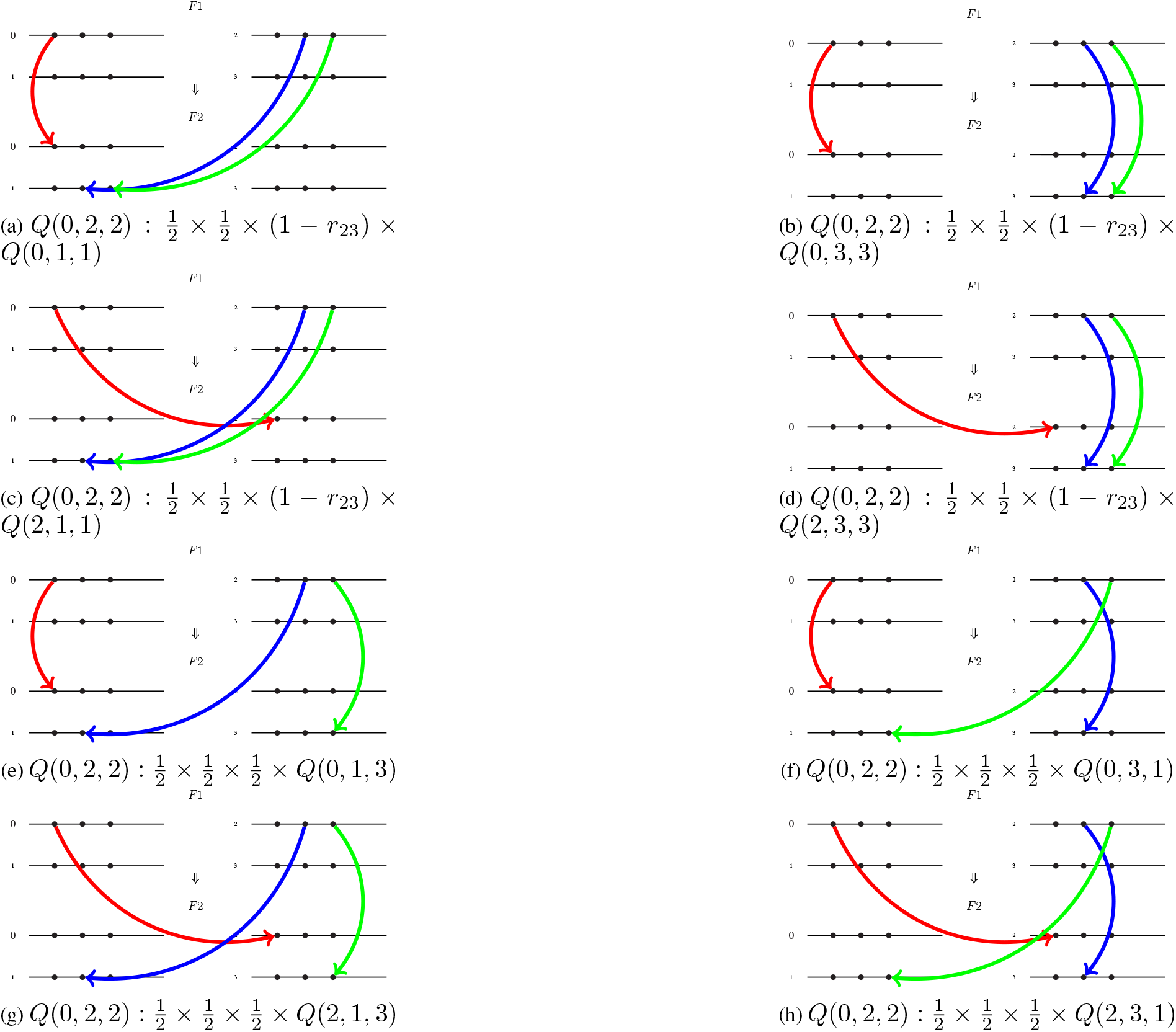
The graphical representation of the factors multiplying each *Q* on the right-hand side of Eq. S14 for *Q*(0, 2, 2).

### 2.10 The self consistent equation for *Q*(0, 2, 3)

Figure S10 displays the 8 factors in the self-consistent equation for *Q*(0, 2, 3):

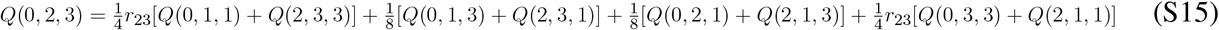

After use of symmetry to keep only non-equivalent *Q*s, this leads to

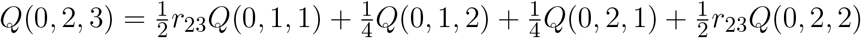

**Figure S10:**
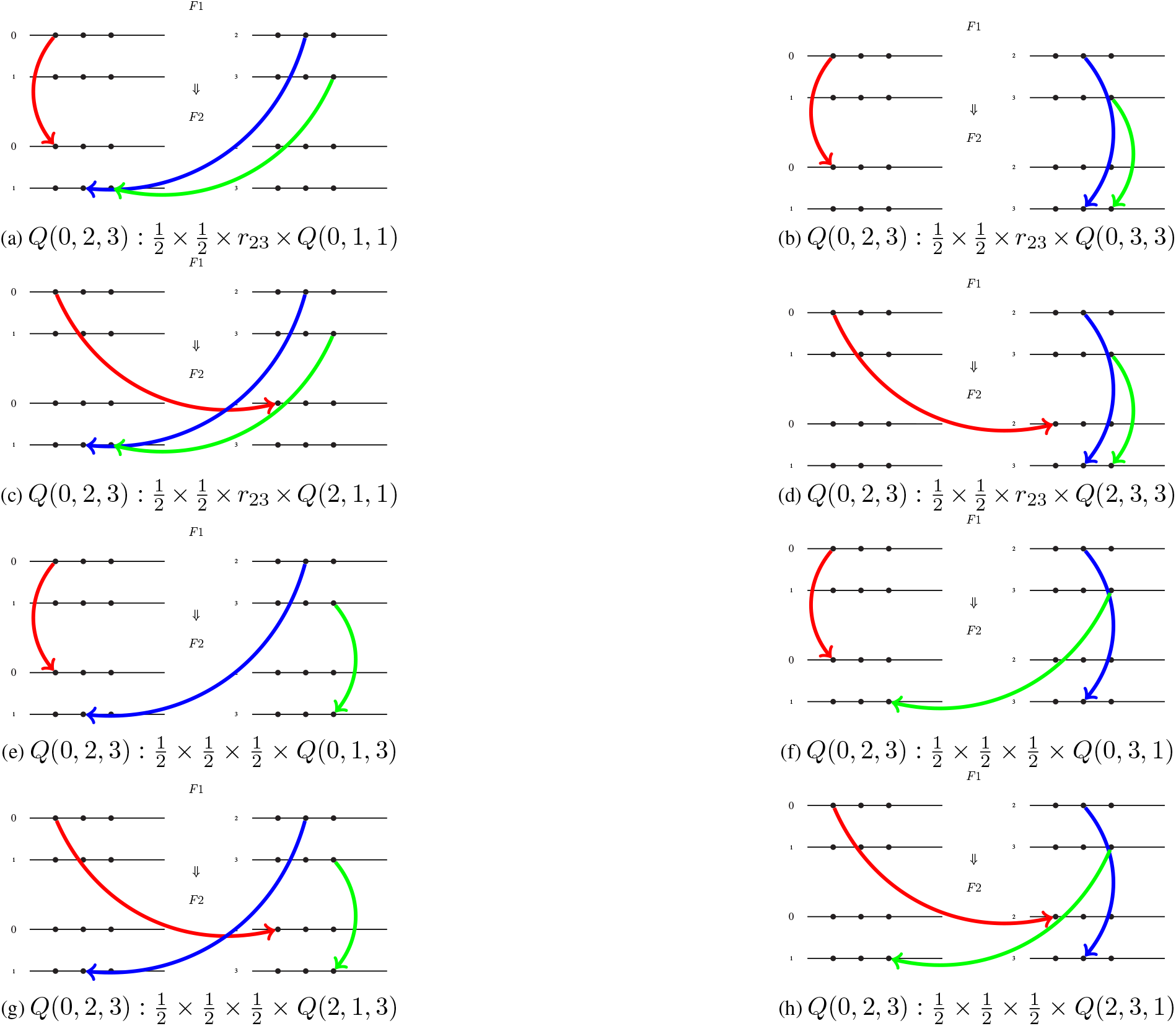
The graphical representation of the factors multiplying each *Q* on the right-hand side of Eq. S15 for *Q*(0, 2, 3).

